# A computer vision-based approach for estimating carbon fluxes from sinking particles in the ocean

**DOI:** 10.1101/2024.07.06.602339

**Authors:** Vinícius J. Amaral, Colleen A. Durkin

## Abstract

The gravitational settling of organic particles in the ocean drives long term sequestration of carbon from surface waters to the deep ocean. Quantifying the magnitude of carbon sequestration flux at high spatiotemporal resolution is critical for monitoring the ocean’s ability to sequester carbon as ecological conditions change. Here, we propose a computer vision-based method for classifying images of sinking marine particles and using allometric relationships to estimate the amount of carbon that the particles transport to the deep ocean. We show that our method reduces the amount of time required by a human image annotator by at least 90% while producing ecologically- informed estimates of carbon flux that are comparable to estimates based on purely human review and chemical bulk carbon measurements. This method utilizes a human-in-the-loop domain adaptation approach to leverage images collected from previous sampling campaigns in classifying images from novel campaigns in the future. If used in conjunction with autonomous imaging platforms deployed throughout the world’s oceans, this method has the potential to provide estimates of carbon sequestration fluxes at high spatiotemporal resolution while facilitating an understanding of the ecological pathways that are most important in driving these fluxes.

## Introduction

The ocean is responsible for regulating the amount of carbon dioxide (CO_2_) that persists in the atmosphere. The difference in partial pressure of CO_2_ across the air–sea interface drives dissolution and fixation of CO_2_ into organic biomass by phytosynthetic algae in surface waters. A fraction of this biomass is packaged into particles and sinks down the water column as particulate organic carbon (POC), where the carbon is stored over long timescales (Ducklow et al., 2001; Boyd et al., 2019). Thus, accurately constraining POC export is import for quantifying the ocean’s role in removing carbon dioxide from the atmosphere.

Technological advances in recent years have facilitated widespread collection of imaging data from the ocean, which presents an opportunity for estimating carbon fluxes with high spatiotemporal resolution (Lombard et al., 2019; Giering et al., 2020). For example, the Underwater Vision Profiler (UVP; Picheral et al., 2010) has been used to image particles in situ and estimate the fluxes that they contribute based on the sizes of observed particles (Clements et al., 2022, 2023). However, uncertainties in UVP-based flux estimates can exceed 50% (Bisson et al., 2022), likely because particles are typically considered monolithically, with a uniform relationship to carbon content and sinking speed. In actuality, the particles responsible for carbon export are highly diverse, being formed by a variety of ecological and physical processes that in turn alter their carbon content and sinking speeds.

Durkin et al. (2021) showed that ecological classification of particles enables relatively accurate estimates of carbon export. However, this approach relied on manual annotation of images for all particles considered in the flux calculations, which is extremely costly and does not scale to large datasets. Trudnowska et al. (2021) used an unsupervised (i.e., not requiring manual annotation) approach based on principal component analysis to categorize particles imaged in the water column by the UVP. This approach has the advantage of removing human bias from categorization, but introduces ambiguity into translating statistical categories into distinct classes of known ecological source and theoretical carbon content.

Convolutional neural networks (CNNs) are commonly used for the task of image classification, and have been applied in the aquatic environment to identify species of phytoplankton (Orenstein and Beijbom, 2017; Cheng et al., 2019; Guo et al., 2021) and zooplankton (Dai et al., 2016; Hong et al., 2020; Li et al., 2021). These CNNs are usually trained with a supervised learning approach, in which an expert manually labels a subset of images from a given sampling campaign that are used for training. The resulting CNN is then used to predict labels from other regions or time periods (i.e., other “domains”). However, there is an implicit assumption that the target domain distribution (i.e., the data that the CNN is used to predict on) should match the distribution of the training domain (Daume III and Marcu, 2006). This is rarely applicable in the dynamic marine environment, where phytoplankton and zooplankton community structure varies greatly with space and time, resulting in distribution shift (Orenstein et al., 2020). Domain adaptation, which refers to the inclusion of data from the target domain in the training set, may aid in mitigating CNN performance degradation due to distribution shift (Kay et al., 2022).

CNNs have also been applied in semi-supervised approaches, which require the human annotator to review only a fraction of imaged particles while clustering similar images together (Schröder et al., 2020; Schröder and Kiko, 2022). This approach has the potential to reduce the subjectivity of a human annotator, but its success depends on how well the clustering algorithm can assign images to ecologically important categories. Particles left unclassified may take a significant amount of time to review.

In this paper, we propose a novel CNN-based methodology for classifying imaged particles that allows us to model particle carbon content with more granularity than with size alone, and may lead to more accurate predictions of carbon fluxes while diagnosing which ecological pathways contribute most to these fluxes. Our method utilizes a human-in-the-loop domain adaptation approach to address the dataset shift problem and to facilitate data assimilation from future sampling campaigns. We use allometric relationships to quantify the carbon content in labeled particles, and compare the resulting flux estimates to those from other more traditional methods of estimating carbon fluxes. Here we apply this approach to microscopy images of particles collected in sediment traps, but the general methodology could be applied to the classification of any particle imaging instrument. If combined with autonomous particle imaging platforms, this method would allow for estimation of carbon fluxes at high spatiotemporal resolution and facilitate an understanding of how the magnitude of carbon export is changing throughout the world’s oceans.

## Materials and procedures

### Data

#### Sampling locations

Particle samples were obtained from the central and subarctic North Pacific, the Santa Barbara Basin, and the North Atlantic (Figure 1). In the central North Pacific, three stations were sampled between Hawai‘i and California aboard the R/V Falkor between January 24 and February 20, 2017. These stations included oligotrophic low flux regions in the subtropical North Pacific, as well as a coastal environment in the California Current (measured POC flux: 1.1–1.7 mmol C m*^−^*^2^ d*^−^*^1^) (Durkin et al., 2021, see their Table 1). Samples from the subarctic North Pacific come from first the NASA EXPORTS field campaign, which took place near Station P between August 14 and September 9, 2018 aboard the R/V Roger Revelle (Siegel et al., 2021). Station P is a high nutrient low chlorophyll region characterized by low export flux (0.4–2.8 mmol C m*^−^*^2^ d*^−^*^1^). Another station was sampled in the Santa Barbara Basin aboard the R/V Sally Ride between December 12–17, 2019, where the settling flux of POC from surface waters was relatively high (5.0– 6.6 mmol C m*^−^*^2^ d*^−^*^1^). Finally, samples from the eastern North Atlantic were collected aboard the R.R.S. James Cook between May 6–24, 2021 during the second NASA EXPORTS field campaign near the Porcupine Abyssal Plain (Johnson et al., 2024). Sampling was conducted in a mesoscale eddy during the spring bloom, which was a high flux system (2.1–11.2 mmol C m*^−^*^2^ d*^−^*^1^).

**Figure 1:**
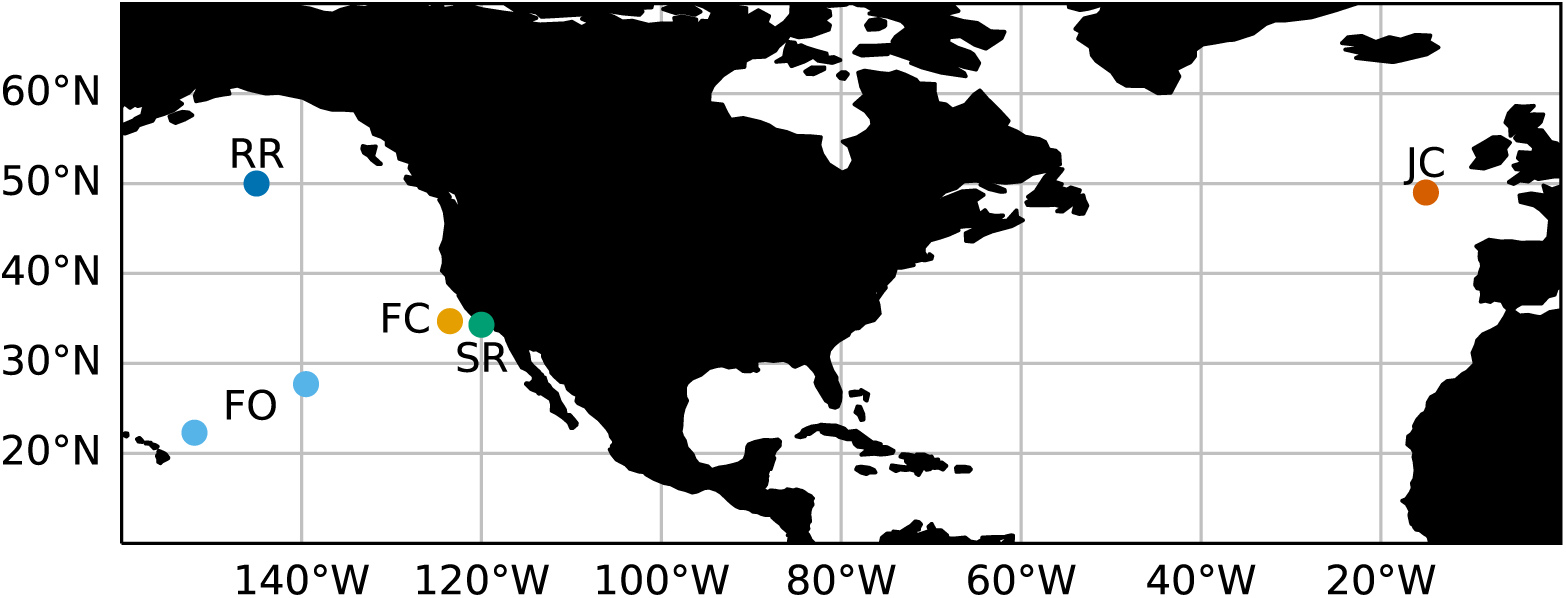
Map of sampling locations, including the subarctic North Pacific (RR), central North Pacific (FO), California Current (FC), Santa Barbara Basin (SR), and North Atlantic (JC).

**Figure 2:**
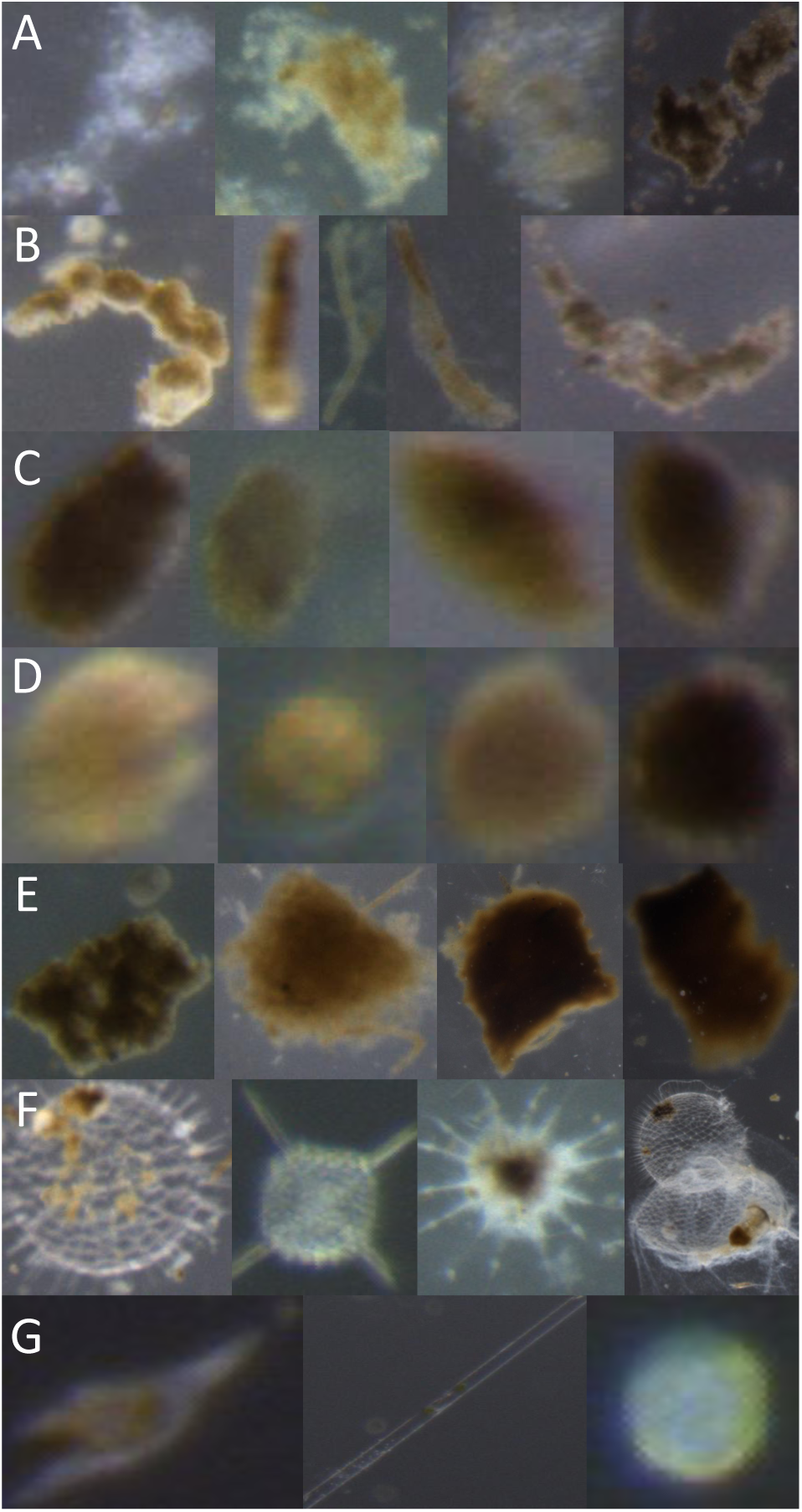
Particles types that are considered for flux calculations including (A) aggregates, (B) long pellets, (C) short pellets, (D) mini pellets, (E) salp pellets, (F) rhizaria, and (G) phytoplankton, including (from left to right) an example of a dinoflagellate, a “long,” and a “round” phytoplankter. Images are not to scale.

**Table 1.** Equations and parameters used to model the carbon content of each particle class.

| Class | Shape | Width | Length | Volume | $A$ | $B$ | Ref |
| --- | --- | --- | --- | --- | --- | --- | --- |
| aggregate | sphere | $w = \text{ESD}$ | $l = \text{ESD}$ | $V = \frac{4}{3} \cdot \pi \cdot \left(\frac{\text{ESD}}{2}\right)^3$ | $1.13 \times 10^{-10}$ | 0.81 | 1 |
| long pellet | cylinder | $w = \frac{264 \cdot \text{ESD}}{\text{ESD} + 584}$ | $l = \frac{\pi}{w} \cdot \left(\frac{\text{ESD}}{2}\right)^2$ | $V = l \cdot \pi \cdot \left(\frac{w}{2}\right)^2$ | $1.13 \times 10^{-10}$ | 1 | 1 |
| short pellet | ellipsoid | $w = 0.54 \cdot \text{ESD}$ | $l = \frac{\text{ESD}^2}{w}$ | $V = \frac{4}{3} \cdot \frac{l}{2} \cdot \pi \cdot \left(\frac{w}{2}\right)^2$ | $1.13 \times 10^{-10}$ | 1 | 1 |
| mini pellet | sphere | $w = \text{ESD}$ | $l = \text{ESD}$ | $V = \frac{4}{3} \cdot \pi \cdot \left(\frac{\text{ESD}}{2}\right)^3$ | $1.13 \times 10^{-10}$ | 1 | 1 |
| salp pellet | cuboid | $w = 0.63 \cdot \text{ESD}$ | $l = \frac{\pi}{w} \cdot \left(\frac{\text{ESD}}{2}\right)^2$ | $V = l \cdot w \cdot \frac{w}{4}$ | $4 \times 10^{-11}$ | 1 | 2, 3 |
| rhizaria | sphere | $w = \text{ESD}$ | $l = \text{ESD}$ | $V = \frac{4}{3} \cdot \pi \cdot \left(\frac{\text{ESD}}{2}\right)^3$ | $4 \times 10^{-12}$ | 0.939 | 4, 5 |
| phytoplankton | sphere | $w = \text{ESD}$ | $l = \text{ESD}$ | $V = \frac{4}{3} \cdot \pi \cdot \left(\frac{\text{ESD}}{2}\right)^3$ | $2.88 \times 10^{-10}$ | 0.811 | 4 |

Parameters $A$ and $B$ in Equation 1 come from various reference (Ref) studies: (1) this study, (2) Silver and Bruland (1981), (3) Iversen et al. (2017), (4) Menden-Deuer and Lessard (2000), and (5) Stukel et al. (2018). ESD = equivalent spherical diameter.

For the purpose of this study, each sampling campaign will constitute a “domain,” i.e., a region characterized by a unique distribution of sinking particles that was sampled during a given time interval. Each domain is hereafter referred to by an abbreviation given by the vessel that was used for sampling: FO and FC for the oligotrophic and coastal central North Pacific, respectively (sampled aboard the R/V Falkor), RR for the subarctic North Pacific (sampled aboard the R/V Roger Revelle), SR for the Santa Barbara Basin (sampled aboard the R/V Sally Ride), and JC for the eastern North Atlantic (sampled aboard the R.R.S. James Cook).

#### Sample collection

Particle samples were collected as described in Durkin et al. (2021). Briefly, sediment traps were fitted with collection tubes containing a jar with a polyacrylamide gel layer overlaid by filtered seawater (Durkin et al., 2015). Following trap recovery, the tubes were allowed to sit for roughly one hour before water was carefully pipetted off. Micrographs of gel layers were imaged on a stereomicroscope under oblique illumination. Regions of interest (ROIs) that contained individual particles were extracted from each micrograph with an imaging processing protocol described by Durkin et al. (2021). This imaging protocol also generated measurements of equivalent spherical diameter (ESD) of each particle.

#### Data labeling

We classified ROIs based on the ecological provenance of the particles (Figure 2). Our definitions were modified from Durkin et al. (2021), and are summarized here. Aggregates are detrital particles with irregular edges that (i) may have formed from processes such as the physical coalescence of algal cells, or (ii) may be highly-degraded fecal material. Long pellets are fecal pellets that are produced by zooplankton such as euphasiids. Fecal pellets that are relatively short or ovular in shape, such as those produced by larvaceans, were classified as short pellets. Mini pellets are smaller, approximately spherical fecal pellets that are likely produced by smaller organisms such as rhizaria and other microzooplankton. While all other particle types consist of detrital material, individual organisms that sinking passively may also contribute to downward carbon flux. In our samples, such “particles” include rhizaria and phytoplankton. Phytoplankton were separated into dinoflagellates, and “long” (e.g., pennate diatoms) and “round” (e.g., centric diatoms) groups. There are also some classes of ROIs that contain particles that do not contribute to POC export, but that were common enough in our dataset to warrant identification so as to not be counted towards the particle flux. These include zooplankton that likely swam into the trap, fibers (either synthetic or naturally occurring), bubbles (pockets of air trapped in the gel), and noise (empty ROIs that were artifacts of the image processing procedure).

Prior to this study, we manually classified all images from the RR and JC domains. We noticed that many images were “ambiguous,” meaning that they could not definitively be given a unique label out of the set of particle classes enumerated above, because (i) they could justifiably be described by at least two labels, (ii) they were unidentifiable (e.g., too blurry) and/or (iii) they could not be described by any of the particle classes (e.g., consider a fragment of plastic sinking through the water column, but note that these were extremely rare and did not warrant the creation of a separate class). In order to quantify this ambiguity, we relabeled subsets of roughly 3000 and 6000 images from the RR and JC datasets, respectively. These domains were chosen because all images from these domains were annotated by a human, while some images from other domains were not. We observed that roughly 81% and 78% of the new labels matched the original annotations for RR and JC, respectively. Thus, we chose a conservatively defined subset of unambiguously labeled images from each domain to train the models, yielding the following image counts for each domain: (RR) 30300 images, 9078 labeled; (FC) 5454 images, 1186 labeled; (FO) 1799 images, 353 labeled; (SR) 16522 images, 4091 labeled; (JC) 115368 images, 35274 labeled (Figure 3).

**Figure 3:**
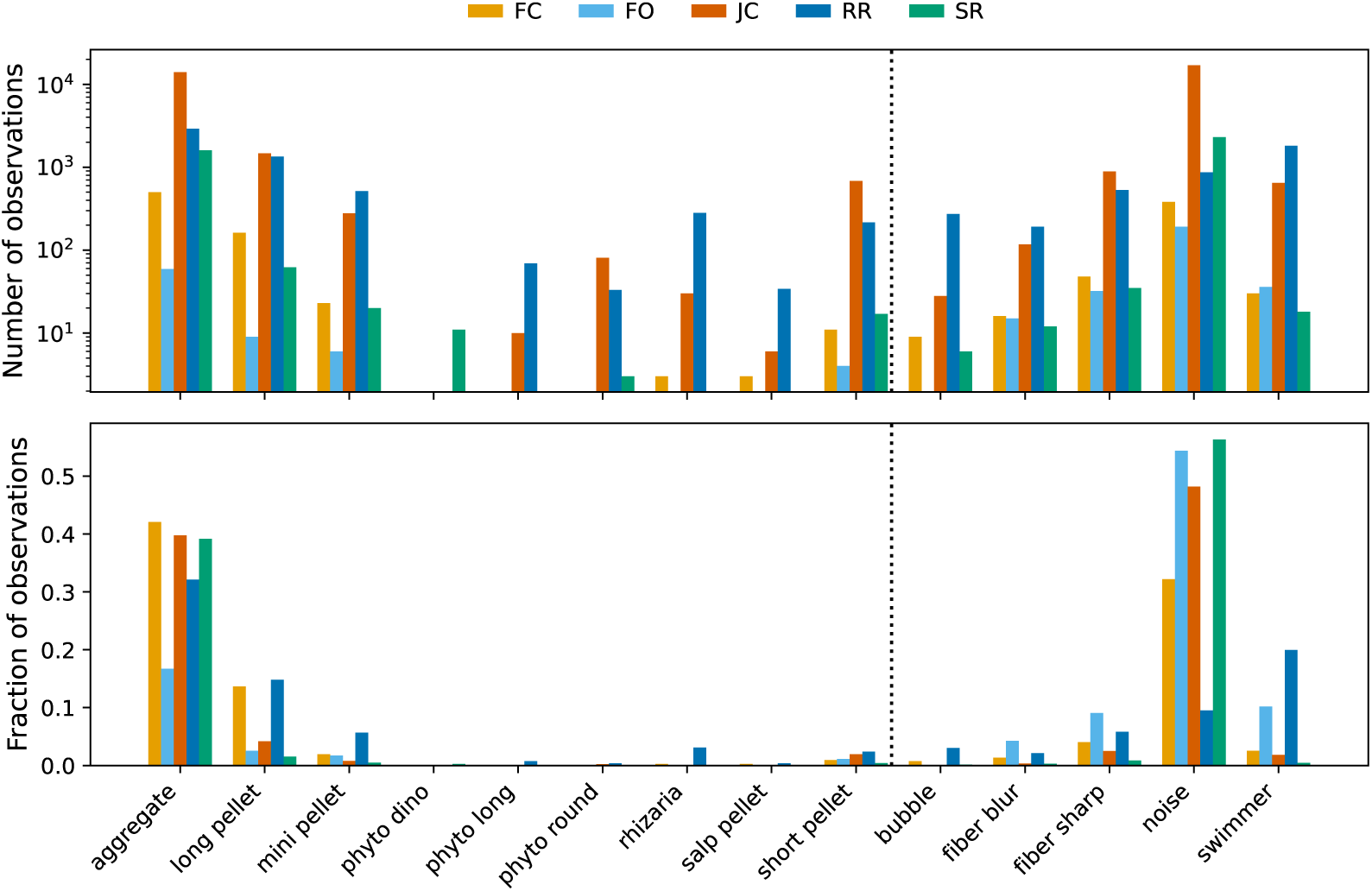
Distribution of labeled particles by class. Classes to the left of the dotted lines are used for the domain adaptation experiments.

Finally, note that in the original set of human-annotated labels that were previously obtained, phytoplankton were not separated into the subclasses described above, noise and bubbles were grouped as “unidentifiable,” and whereas we separated fibers in visually distinct “sharp” and “blur” types, the original labeling scheme did not. We maintained our revised labels (with finer categorization) for CNN training and predictions, but in comparisons to original labels presented later on in this work, our labels were grouped as consistent with the original labeling scheme.

### Hyperparameter tuning

For CNN (hereafter, “model”) training, we selected the ResNet-18 neural network architecture (He et al., 2016) due to its balance between training speed and accuracy (Canziani et al., 2017). Following Orenstein and Beijbom (2017), we finetuned models that were pre-trained on roughly one million images spanning one thousand object classes from the natural and built environments (ImageNet; Russakovsky et al., 2015). Roughly 95% of our images had a longest dimension (i.e., width or height) that was shorter than 224 pixels, so we set the input size to this value in order to minimize obfuscation of particle morphologies via image shrinking. Training was done in epochs, where one epoch describes one pass of the entire training and validation sets through the model. Images were passed through the model in batches of 128, and were shuffled into new batches between epochs. Early stopping of training was implemented with a patience of 10, such that training stopped after there were 10 consecutive epochs without improvement relative to the lowest validation loss. The optimizer (i.e., algorithm used to fit model parameters to the training data by minimizing a loss function) that we used was Adam with weight decay (AdamW; Loshchilov and Hutter, 2019). For data augmentation, 90*^◦^* rotations and horizontal and vertical flips were applied randomly to the images during training. Given this training protocol, we tuned (i) image resizing and normalization, (ii) initial learning rate, and (iii) weight decay by using class-specific precision and recall as evaluation metrics. For each of these hyperparameter tuning experiments, five model replicates were trained with random number generator (RNG) seeds of 0, 1, 2, 3, and 4 to quantify model variance due to RNG initialization. Here, labeled images from domains FC, FO, JC, and SR were used for training and validation while images from RR were used for evaluation (i.e., testing). The train and validation splits were stratified by class, such that for each domain, 80% and 20% of the images were used for training and validation, respectively. All training was done on a NVIDIA RTX 8000 running CUDA 11.6.

First, we investigated the effects of two image resizing techniques and image normalization. ResNet-18 requires square images as input. However, our particle images were usually rectangular and it may be important to preserve their aspect ratio such that one dimension is not scaled without a proportional scaling of the other (e.g., a short pellet that is stretched only along its shorter axis may resemble a mini pellet). To resolve this issue, we centered images between black borders (i.e., zero-padding). Images that had a longer dimension greater than 224 pixels were shrunk while preserving aspect ratio, and black borders were added on either side of the image along the shorter dimension. Images with a longer dimension that was less than 224 pixels were simply zero-padded (Hashemi, 2019). This preprocessing protocol, referred to herein as “CustomPad,” was compared to Resize from PyTorch’s torchvision.transforms module, which simply resizes both image dimensions to 224 with no aspect ratio preservation.

In addition to image resizing, we also evaluated how data normalization affected our evaluation metrics. The mean and standard deviation calculated from the RGB channels of ImageNet images ([0.485, 0.456, 0.406] and [0.229, 0.224, 0.225], respectively) are commonly used for data normalization. The mean and standard deviation calculated from our training dataset after applying CustomPad were [0.053, 0.058, 0.055] and [0.123, 0.133, 0.127], respectively. Using Resize on the other hand, yielded a mean and standard deviation of [0.279, 0.304, 0.294] and [0.096, 0.102, 0.095], respectively. To quantify model sensitivity to image resizing and data normalization, we trained models with 6 combinations of resizing and data normalization protocols: (i) Resize with no normalization, (ii) CustomPad with no normalization, (iii) Resize with normalization via statistics calculated from our Resize-transformed data, (iv) Resize with normalization via ImageNet statistics, (v) CustomPad with normalization via statistics calculated from our CustomPad-transformed data, and (iv) CustomPad with normalization via ImageNet statistics. For these experiments, initial learning rate and weight decay were fixed to the AdamW defaults of 0.001 and 0.01, respectively. We found no sensitivity to image resizing and data normalization based on our evaluation metrics (Supplemental Figure S1), thus we proceed with the simplest protocol of resizing with Resize and no normalization.

Next, we fixed weight decay at 0.01 and varied the initial learning rate across three orders of magnitude: 0.0001, 0.001, and 0.01. We found that compared to the default value of 0.001, the higher initial learning rate degraded performance as measured by our evaluation metrics, while the lower learning rate did not noticeably affect performance (Supplemental Figure S2). Thus, we maintained the default learning rate of 0.001.

Finally, we tuned weight decay by considering three orders of magnitude for this parameter as well: 0.001, 0.01, and 0.1. In our experiments, the choice of weight decay did not affect model performance (Supplemental Figure S3), so we maintained the default value of 0.01. All model training subsequently described in this study was thus done with image resizing that does not preserve aspect ratio (i.e., Resize), no image data normalization, an initial learning rate of 0.001, and weight decay set to 0.01.

### Domain adaptation experiments

Upon obtaining images from a sampling campaign at a novel target domain, we would like to train a model to classify the images with high accuracy while minimizing human involvement. Ideally, the distribution used to train a model should be the same as that which is being classified, i.e., the target set (Daume III and Marcu, 2006). In reality, this approach is often impossible to apply if the underlying distribution of a novel unlabeled set of particles is unknown. Furthermore, the particle morphologies for a given class may vary from region to region, e.g., an aggregate from one domain may look different than an aggregate from another domain. One approach may be to manually label a subset of images from each novel sampling campaign in order to finetune a model, but this approach does not scale to large datasets because (i) it is not clear how many images an expert must annotate in order to capture the true distribution of the dataset and (ii) obtaining such labels is expensive. Although intra-class morphological variance between domains may exist, feature representations learned in one domain may transfer to a separate target domain.

In order to take advantage of knowledge gained from labeled data from previous sampling campaigns while minimizing human effort, we propose a human-in-the-loop domain adaptation (Zhou et al., 2022) approach in which first, an out-of-domain (OOD) model ensemble is finetuned with images labeled from previous sampling campaigns. This OOD ensemble is then used to predict labels for the novel domain, and ensemble voting is used to propose images to show to a human expert for verification. Next, the in-domain verified images are added to the previously OOD training set, and the ensemble is finetuned once again and used to predict labels for all remaining in-domain images not included in the training set. Finally, carbon fluxes are calculated from the combination of human-verified and model-predicted labels. Our methodology is summarized in Figure 4 and elaborated below.

**Figure 4:**
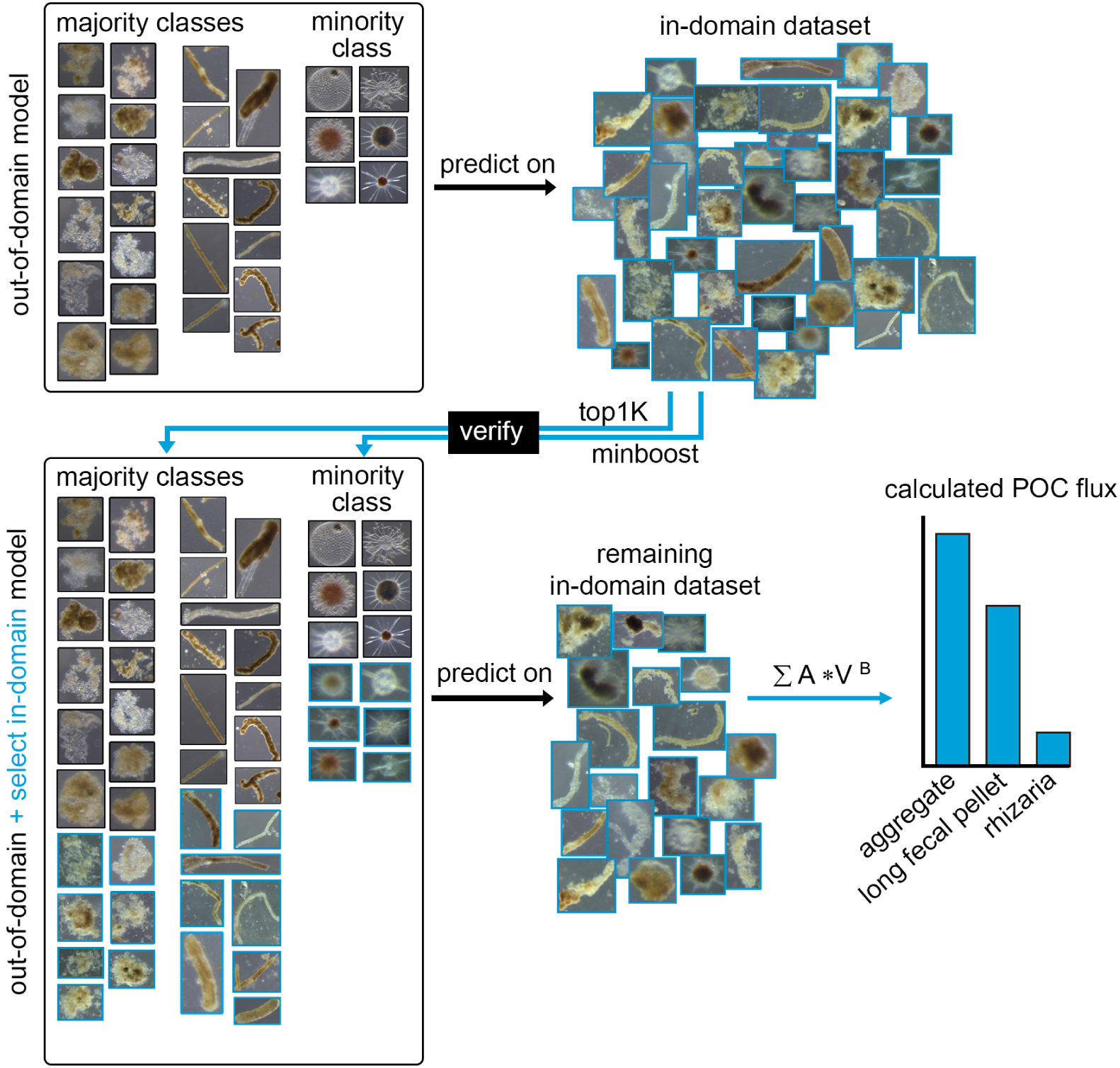
Summary of the entire human-in-the loop classification method. First, a model is trained on out-of-domain images. This model is used to predict labels for in-domain images, a subset of which are verified by a human expert. The verified images are combined with the original out-of-domain images to train another model, which is used to classify all remaining in-domain images not included in the training set. Finally, fluxes are calculated for each particle class.

#### Out-of-domain model training

Our methodology was tested with both RR (subarctic North Pacific) and JC (North Atlantic) as the target domain. This is because for both domains, we previously labeled all particle images manually and thus have a human baseline with which to compare our model-based labels and flux calculations. We also re-labeled a subset of the images from each of these two domains to quantify intra-annotator variance that may be caused by ambiguity in particle morphologies. For a given target domain, training and validation sets were compiled from all other domains and an OOD model ensemble was finetuned as described for hyperparameter tuning.

#### Model ensemble voting

After the OOD ensemble was trained, each ensemble replicate was used to predict confidence scores corresponding to each particle class for every image in the target domain. The Softmax function (torch.nn.Softmax) was used to transform ResNet-18’s output vector of logits into a vector of confidence scores between 0 and 1 (we emphasize that these scores should not be interpreted as probabilities, see Guo et al., 2017), and the particle class with the highest score was taken as the image label. For all images that had unanimous label consensus across all five ensemble replicates, the mean score for the consensus label across all replicates was calculated. The images with the 1000 highest mean scores for each class were shown to a human expert for verification. Note that some classes may have had fewer than 1000 images with unanimous consensus between the ensemble replicates, indicating that the expert had fewer than 1000 images to review for these classes.

#### Human verification

The suggested images from the model ensemble voting step were organized into a directory with subdirectories named by particle class. The expert verified the images by reviewing the images in each class directory. If the image was labeled incorrectly and there was no ambiguity as to what the correct label should have been, the image label was corrected by moving the image to the subdirectory corresponding to the correct label. If there was ambiguity regarding the label of a suggested image, then the image was simply deleted. Otherwise, if the label was correct, no action was taken.

#### Finding more minority class instances

Minority classes in the OOD training set may be poorly learned, resulting in few or no consensus instances suggested by the unanimous voting scheme. In the directory of images verified by the expert, any classes containing fewer than 100 instances were considered to be minority classes. For each of these classes, the images whose scores appeared in the top 1000 scores across all replicates were suggested for verification in a new directory whose subdirectories were named by minority class. The expert simply deleted images that were incorrectly labeled. The suggested images in this step did not include images that were manually verified in the previous step as a result of unanimous consensus among the OOD model ensemble. Note that a class may have been relatively abundant in the OOD set but may still have had fewer than 100 instances in the in-domain suggested set, thus being considered a minority class in this step.

#### Model retraining

The manually verified in-domain images were incorporated into new training and validation sets. These images were split 80%/20% and stratified by class. The 80% subset was combined with all OOD images (used in both the training and validation sets for the OOD ensemble) to form the new training set. The validation set was composed only of the 20% split of verified indomain images in order to fit the model only to target domain data. A model ensemble was then finetuned as before, using ImageNet weights as a starting point. This ensemble was used to predict labels for all remaining in-domain images, i.e., those that were not integrated into the training and validation sets.

### Carbon flux estimates

Once all particles from the target domain were labeled, POC fluxes were calculated for each gel trap, similar to Durkin et al. (2021) with slight modification to some parameters. We updated the parameters used to model POC fluxes because we combined classes that were previously split into separate categories and because more measured POC flux data was available to fit model parameters. Here, we parameterize a single “aggregate” category (previously split into aggregates and dense detritus) and a single long fecal pellet category (previously split into long fecal pellets and large-loose fecal pellets), in addition to the five other particles contributing to POC flux (see Table 1). Combining the previous nine categories into seven reduced the inconsistency in both the human and machine classification of the most visually diverse and sometimes ambiguous particle classes.

The mass of carbon *C* (mg) of a single particle is given by

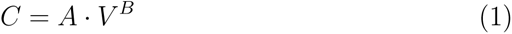

where *A* is a scaling coefficient (mg µm*^−^*^3^, essentially the carbon mass of 1 µm^3^), *V* is the volume of the particle (µm^3^), and *B* is an exponent parameter (unitless) that describes carbon density. The volume *V* is modeled to best approximate the shape of each particle type and is a function of the equivalent spherical diameter (ESD, µm) of the particle. For particles whose volumes were approximated as spherical (aggregates, mini pellets, rhizaria, phytoplankton), the ESD was used to estimate the radius of the sphere to calculate *V*. The volumes of other particle types were estimated as cylinders (long fecal pellets), ellipsoids (short fecal pellets), or cuboids (salp fecal pellets), requiring length and width measurements not accurately estimated by automated image processing functions. Durkin et al. (2021) measured the width of 186 salp fecal pellets, 596 short fecal pellets, 563 large-loose fecal Table 1: Equations and parameters used to model the carbon content of each particle class pellets, and 1415 long fecal pellets to identify an empirical relationship with ESD calculated from measured particle area. Here, we use these previously published parameters relating ESD to width for salp fecal pellets and short pellets, best approximated by a linear relationship. Because we combined the long fecal pellet and the large-loose pellet categories, we identified a new combined relationship relating ESD to pellet width for this category, which is best described by a hyperbolic relationship (Durkin et al., 2021, see their Table 1). Lengths of cylinders, ellipsoids, and cuboids were then described as a function of width and ESD, as described by Durkin et al. (2021).

To convert volumes into carbon units, the *A* and *B* parameters for each particle type were modeled using a minimization function (scipy.optimize.minimize) that gave the best fit to log transformed chemically measured bulk POC fluxes. The same modeled value of *A* was used for aggregates, long, short, and mini pellets. The value of *A* used to describe salp fecal pellets, phytoplankton, and rhizaria were based on literature values (Table 1). The value of *B* was modeled only for aggregates, and fixed at 1 for particles whose carbon content is not known to vary as a function of volume. The *B* value of other particles (phytoplankton and rhizaria) was taken from literature values. We used the same datasets as Durkin et al. (2021) to fit these imaging-based parameters of carbon flux to measured carbon fluxes, and also included 11 additional samples collected during the two sediment trap deployments in the North Atlantic (JC). The updated estimates of *A* and *B* model parameters were similar to those in the previous study and did not noticeably change previously reported results.

After calculating the mass, *C*, of carbon in each particle using Equation 1 and the updated parameters, POC flux was calculated by dividing the mass by the total area imaged for the relevant magnification and the total deployment time for the trap from which the sample originated. Fluxes of each particle category were summed to calculate the total flux in each gel trap, as predicted by each of the model replicates. Thus, variability in flux estimates for a given sample arose from differences in predictions for unverified particles between model replicates. Fluxes were calculated when considering each of RR and JC as the target domain, with 30 and 20 gel trap samples from these domains, respectively.

## Assessment

In order to establish a human baseline against which to compare our modelbased flux calculations, first we calculated fluxes based on the expert annotations and compared those flux estimates to measurements of bulk carbon from the RR and JC datasets presented in Durkin et al. (2021) and Estapa et al. (2021), and Siegel et al. (unpubl.), respectively. We found a mean absolute error (MAE) between the flux estimates from human annotations and those from bulk carbon measurements of 0.71 mmol C m*^−^*^2^ d*^−^*^1^ and 1.55 mmol C m*^−^*^2^ d*^−^*^1^ for the RR and JC datasets, respectively (Figure 5). Next, we calculated fluxes that incorporated model-based predictions of particle classes. In order to examine the effect of each step in our proposed domain adaptation methodology, we considered four sets of predictions for each target domain in order to calculate fluxes. The first set of predictions arose from the OOD model, whose training set only included out-of-domain images. In the second set of predictions (+top1k), flux calculations were based on a model ensemble that was retrained on up to 1000 images from each class that were labeled by the OOD model. Human verification of these images was used in the third set of predictions (+verify). A final ensemble voting technique was applied to improve predictions of minority classes (+minboost).

**Figure 5:**
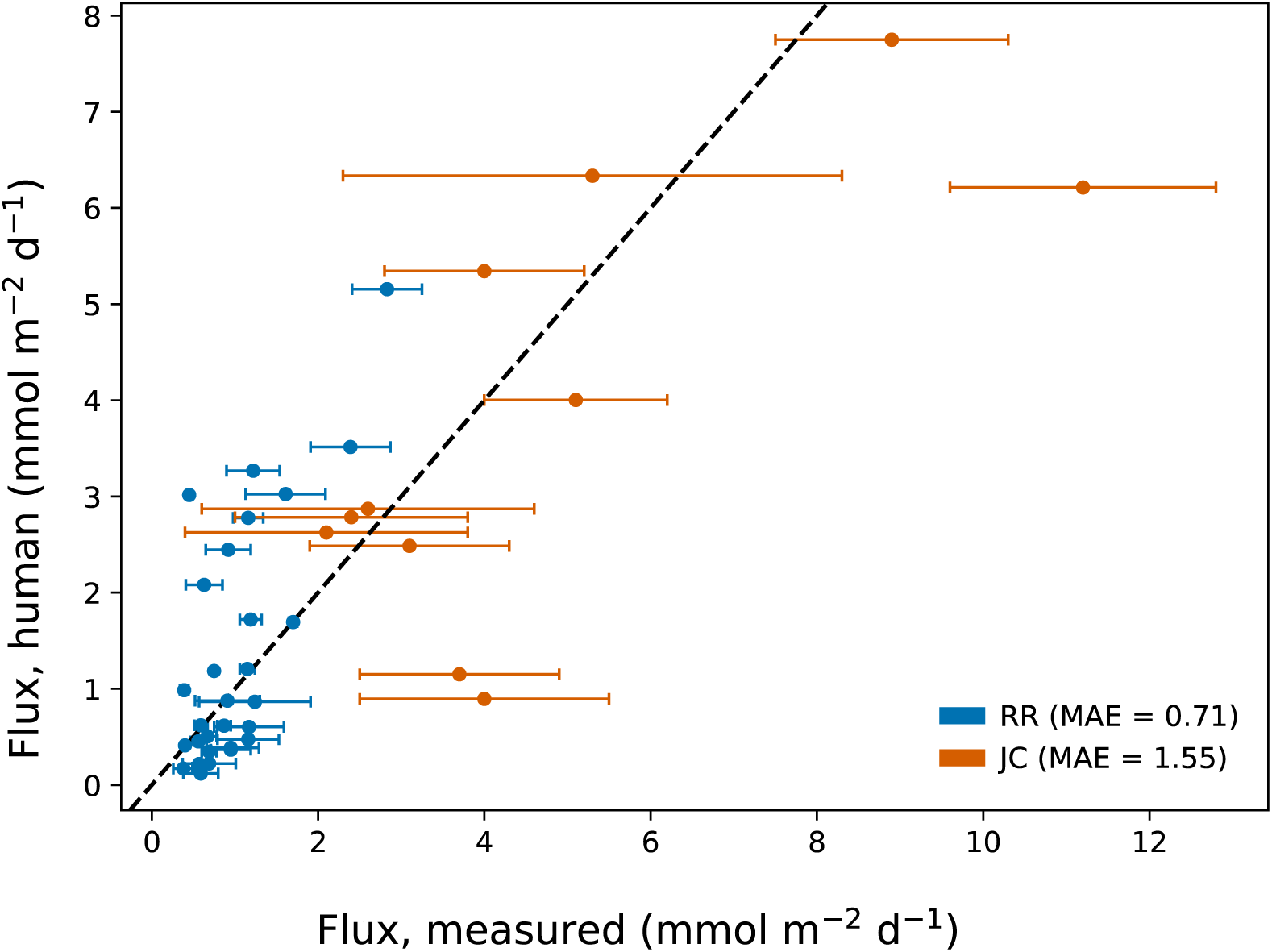
Comparison of fluxes calculated from the original human-annotated labels versus bulk carbon measurements from the traps from the North Pacific (RR) and North Atlantic (JC) sampling campaigns. Each marker represents one sediment trap sample. Dashed lines denote perfect agreement between the different estimates of carbon fluxes. MAE is the mean absolute error. Error bars represent the standard deviation of replicate sample splits (see Durkin et al., 2021).

We compared the MAE from fluxes calculated from model predictions to those calculated from human annotations (total and by class), as well as the MAE between total flux estimates from model predictions and bulk carbon measurements (Figure 6). The variance was generally largest for the flux estimates from OOD predictions relative to those from the domain adaptation refinements. The incremental steps in the domain adaptation experiment appeared to improve (though not monotonically) the MAE between total fluxes estimated from model predictions and both those from human annotations (“total”) and bulk carbon measurements (“measured”). Notably, estimates from the domain adaptation treatments that involved human verification (+verify and +minboost) had MAEs that were comparable to those between estimates from human annotations and bulk carbon measurements (Figure 6, gray lines).

**Figure 6:**
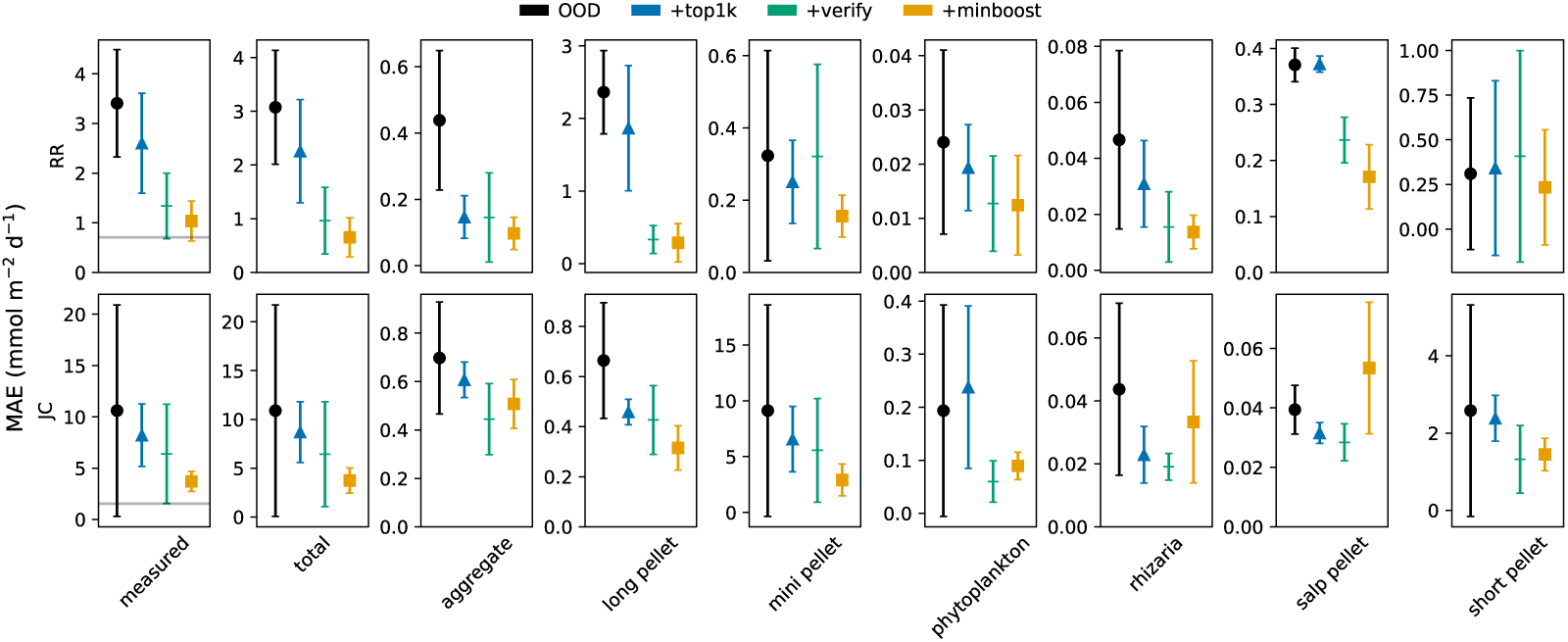
Mean absolute error between fluxes calculated from model labels and (first column) bulk carbon measurements from the traps and (all other columns) the original human-annotated labels, from the North Pacific (RR) and North Atlantic (JC) sampling campaigns. The gray lines in the first column correspond to the MAEs shown in Figure 5. Error bars indicate one standard deviation of flux estimates across five model replicates.

In order to test for differences in significance between model treatments, we conducted analysis of variance (ANOVA) for each panel in Figure 6, followed by a post-hoc Tukey test if ANOVA yielded a significant (*p <* 0.05) result. With RR as the target domain, there were significant improvements in MAE provided by the +verify predictions compared to the OOD predictions for measured and total fluxes, as well as for aggregates, long pellets, and salp pellets. However, +minboost yielded no significant improvement compared to +verify for any total or class-specific fluxes. With JC as the target domain, there was only a significant improvement for long pellets provided by +minboost relative to the OOD predictions. However, +minboost significantly increased MAE compared to +verify for salp pellets (note however, the high variance of +minboost compared to that of +verify).

Examining the flux-specific MAEs is important in measuring performance relative to fluxes, which is an ecologically relevant metric. However, since MAE has the same units as carbon flux, larger, more abundant particles are more likely to have higher MAEs than smaller, less abundant particles. In order to evaluate model performance on each particle class that is independent of carbon content, we show the class-specific precision and recall for the two target domains (Figure 7). In this comparison, the ground truth labels were considered to be those from the original expert annotations of the entire RR and JC datasets obtained prior to this study, which included ambiguous images. Note that in Figure 7, the noise and bubble classes were grouped as “unidentifiable,” as done in the original expert annotations. In order to quantify ambiguity in the original image labels, we randomly selected and relabeled roughly 3000 and 6000 images from the RR and JC datasets, respectively, and plotted the precision and recall relative to the original annotations as gray lines in Figure 7.

**Figure 7:**
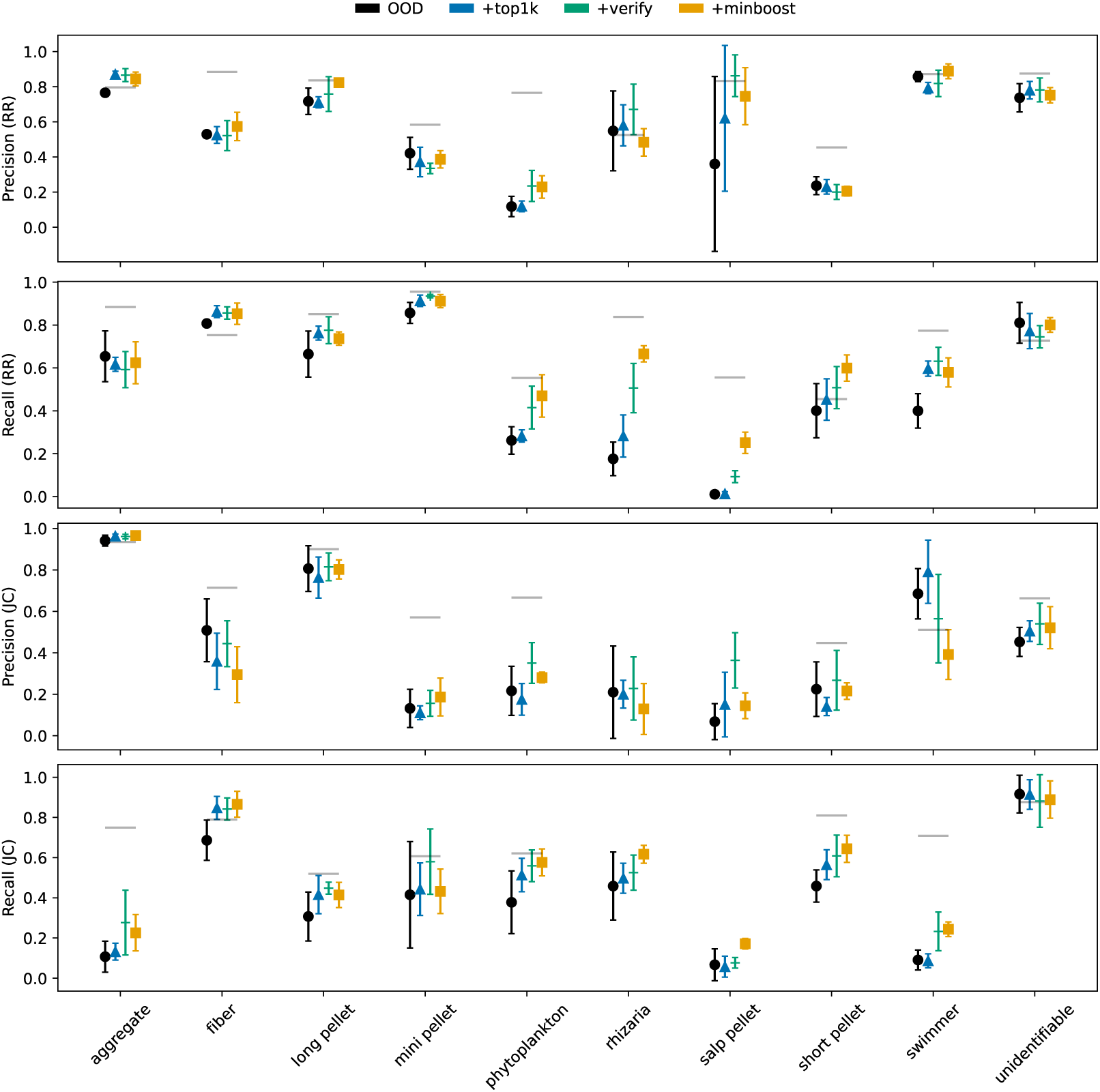
Precision and recall by class and sampling campaign from the domain adaptation experiments. The gray lines show intra-annotator metrics calculated from the relabeling experiments, considering the original manual labels as the ground truth. An absence of gray lines indicates an absence of samples for a given class in the subset of randomly relabeled images. Error bars indicate one standard deviation of flux estimates across five model replicates.

For both target domains, the class-specific precision and recall from the models were often comparable to those from the re-annotation experiment for several classes including aggregates, long pellets, mini pellets, and short pellets. Model performance was noticeably worse relative to the re-annotation metrics for rarer classes such as phytoplankton, rhizaria, and salp pellets. We conducted ANOVA for each domain-metric-class grouping followed by a posthoc Tukey test if ANOVA yielded a significant (*p <* 0.05) result. Compared to the OOD model, +verify significantly improved precision for aggregates (RR), phytoplankton (RR), and salp pellets (JC), as well as recall for mini pellets (RR), phytoplankton (RR), rhizaria (RR), salp pellets (RR), swimmers (RR and JC), fibers (JC), and short pellets (JC). Relative to +verify, +minboost further improved recall for rhizaria (RR) and salp pellets (RR and JC), but worsened precision for JC salp pellets, which may explain the corresponding degradation in MAE observed in Figure 6.

In general, precision and recall for JC were worse than for RR. For all four models, precision for RR aggregates was roughly 0.8, and recall was about 0.6. For JC, precision for aggregates was approximately 0.9 while recall was roughly 0.2, indicating that many aggregates were being classified as other classes. Because aggregates were the most abundant class in JC, misclassifying them as other particle classes may have been responsible for the low precision shown for other classes, such as mini pellets, salp pellets, and short pellets. It is likely that many aggregates were labeled as unidentifiable, as recall of unidentifiables was relatively high (*∼*0.8), while precision was not (*∼*0.8).

Finally, we plotted profiles of fluxes estimated from the +minboost replicates compared to those from human annotation based-estimates and bulk carbon measurements (Figure 8). For most sampling deployments, both the model- and human-based flux estimates approximated the fluxes from bulk carbon measurements. Notably for JC, the model estimates overestimated mini pellet and short pellet fluxes and underestimated aggregate and long pellet fluxes compared to the human estimates. This can be attributed to many particles labeled as aggregates and long pellets by the human to be predicted as mini pellets and short pellets, respectively, by the model.

**Figure 8:**
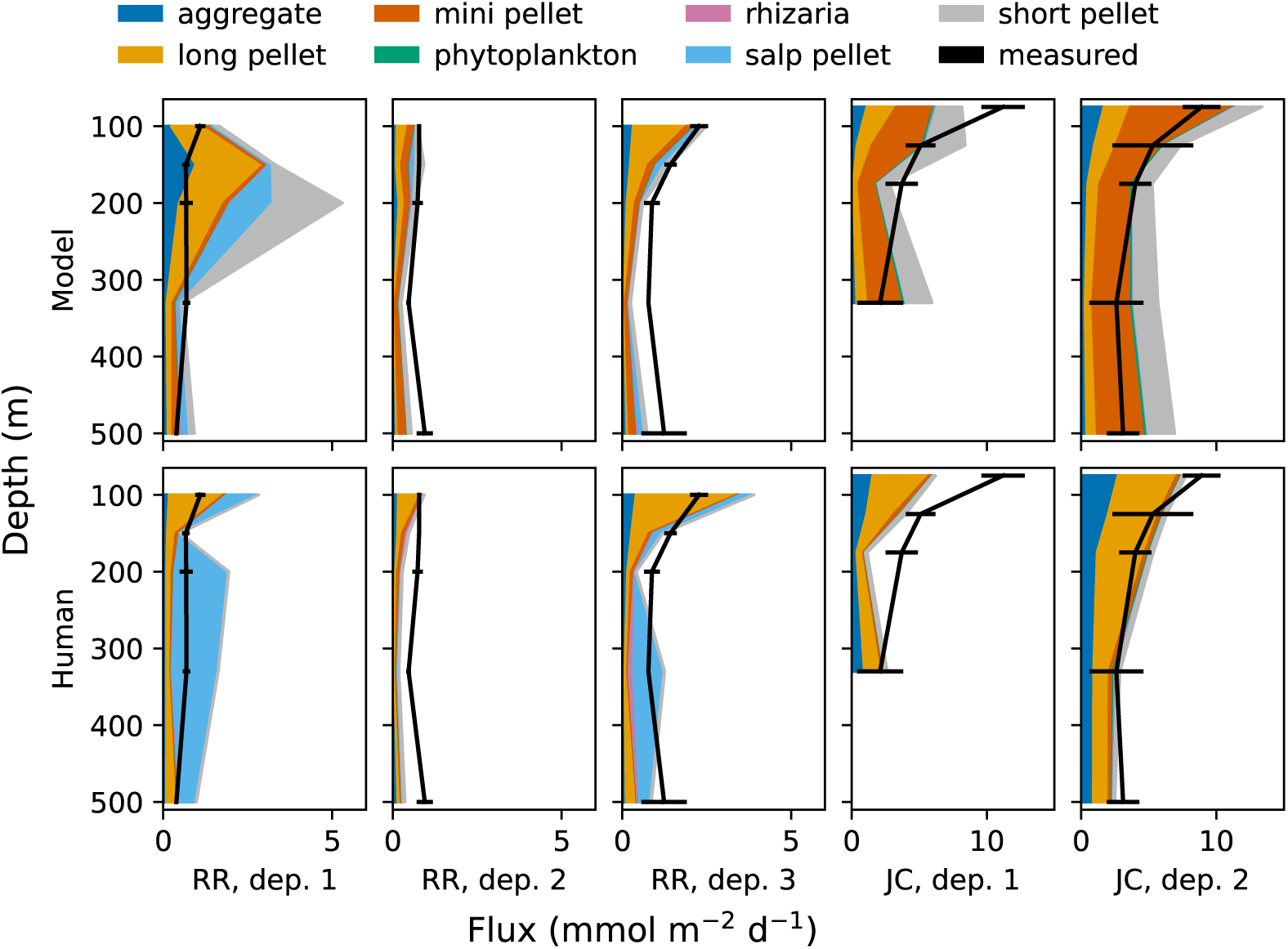
Fluxes estimated from model predictions and human labels from the North Pacific (RR) and North Atlantic (JC) deployments (Dep.) by particle class, as well as from bulk carbon measurements (black). Model estimates are averaged across model replicates. Error bars are propagated from the standard deviation of replicate sample splits at a given depth (see Durkin et al., 2021).

## Discussion

The primary goal of this study was to leverage computer vision to facilitate annotation of particles imaged in the ocean. Manual classification of images from the RR and JC sampling campaigns was done during many months spread out over multiple years. Due to the large number of images that had to be tediously reviewed by (in our case) a single expert, we did not find it feasible to dedicate full workdays over several weeks for this task – shorter intervals over a longer time horizon were critical to maintaining morale and avoiding fatigue. In contrast, verifying labels for up to 1000 images from each class as suggested by the OOD model (+verify) took between roughly 70 (JC) to 90 (RR) minutes, and reviewing additional suggestions from minority classes required about 20 minutes of additional labor (+minboost). Because we did not keep track of the number of hours required for the manual classification done prior to this study, we cannot precisely quantify the savings in human labor. However, we conservatively estimate this figure to be at least 90%.

One clear explanation for the decrease in review time is the reduction in number of images that are reviewed. While all images from a novel sampling campaign must be reviewed in the manual workflow, only up to 1000 images for each class are reviewed in our proposed methodology. A less obvious cause for review time reduction was that in the purely manual approach, not all images were reviewed equally in time. Images that unambiguously belonged to a given class may have been classified in fractions of a second, but image labels that were less clear-cut due to a variety of factors such as visual blurring or morphological ambiguity required more time. The expert annotator may have mulled over ambiguous images for several seconds, and even deferred classification until a future point in the workflow, resulting in a single image being reviewed two or more times. With our method, any image that was reviewed in the first verification step (+verify) had unanimous consensus from the OOD model ensemble. In our experiments, misclassifications were quickly and easily rectified before being integrated into the training set. The second verification step designed to identify more instances of minority classes (+minboost) may have resulted in some images being re-reviewed. This could have occurred, for example, if an image suggested by the OOD model as an aggregate was discarded during the first verification step and was subsequently suggested as a salp pellet in the second step. In our experiments, these instances were rare. Furthermore, while theoretically possible for an image to be suggested as multiple classes in the second step, this was not evident in our experiments. In summary, the entire verification workflow requires an expert to verify only a subset of images from the target domain, most of which are easily and quickly reviewed once.

Not only did our methodology greatly diminish the amount of human effort required to label images, it yielded estimates of total flux that were similar to those calculated from the manual annotations and from imaging- independent estimates based on bulk carbon measurements (Figure 6). One potential net benefit of our approach compared to bulk carbon measurements was that we calculated fluxes contributed by different particle classes, which allowed for diagnosis of which ecological pathways were most relevant for carbon flux (Figure 8). Using class-specific precision and recall as metrics, model classifications performed comparably to human re-annotation for most classes (Figure 7). We propose that the metrics from the human re-annotation experiment are a benchmark for how well we can expect the model to perform. Due to the difficulty of identifying ambiguous images, when using one set of human labels as a ground truth, we should not expect the model to reproduce these labels any better than a human would. Based on this criterium and the similarity between model-, human-, and measurement-based flux estimates described above, we suggest that our method is greatly advantageous in minimizing the amount of human labor required in labeling images, and producing flux estimates comparable to those obtained from human labels and chemical measurements while allowing for diagnosis of prominent ecological pathways in governing carbon flux.

## Comments and Recommendations

Our method is not without its limitations. Consider the low recall for aggregates when JC was the target domain (Figure 7). This is concerning given that aggregates are a majority class in this dataset (Figure 3), and suggests that many aggregates may have been misclassified as other particle types, leading to underestimation of flux for the aggregate class. One hypothesis for inferior performance in JC compared to RR is that the verification steps (+verify and +minboost) resulted in a much smaller proportion of JC images getting integrated into the training and validation sets (4%) compared to RR (21%) for model retraining. This occurred because despite JC having roughly four times as many images as RR, up to 1000 images are reviewed for each class for both target domains. A smaller fraction of the total population integrated into the training set allowed for less data diversity to be learned during training for JC, potentially leading to worse performance.

This issue could be rectified by increasing the number of images to be reviewed by the human expert. Due to the model ensemble voting approach, we expect that the total amount of time required to review additional images would scale linearly with the number of images, given that most of these images were quickly and unambiguously verified in our experiments. We decided to leave the OOD images in the training sets into which in-domain images were incorporated for model retraining. This decision operated under the assumption that particles of a given class look similar enough regardless of what domain they were collected from. In practice, we see that although particles of a given class from two domains shared morphological similarities, they may have been visually distinct (e.g., aggregates from JC were generally less densely packed than those from RR). By increasing the number of images suggested by the OOD model that are then verified by the human expert, we may relinquish the need to maintain OOD images in the model retraining step. Using a purely in-domain training set may lead to better performance for a chosen target domain given that the number of images in this training set is large enough to represent the variance in each particle class.

Finally, we demonstrated that our human-in-the-loop domain adaptation approach (+verify) generally improves classification relative to flux MAE or precision and recall compared to purely OOD predictions. However, the subsequent attempt to boost performance for minority classes (+minboost) has the potential to degrade performance for some particle classes, especially if such classes still suffer from a scarcity of samples after +minboost is applied. We expect that as our method is used to label more and more particles throughout the world’s oceans, feature representations learned by the model for rare classes will improve as these rare samples are added to the training sets, yielding better performance for these classes.

Despite these limitations, we believe that our method is a valuable step in progressing towards an ecologically-informed understanding of carbon flux in the ocean driven by gravitational settling of particles. Compared to statistics-based classification methods (Trudnowska et al., 2021), this approach is based on a categorization scheme derived from pre-defined carbon flux pathways with known ecological significance. Furthermore, like methods developed for similar applications (Schröder et al., 2020; Schröder and Kiko, 2022), our method drastically reduces the amount of human effort required for obtaining classification with the added net benefit that all particles are assigned a label. The human-in-the-loop domain adaptation approach demonstrated here is one that could be applied not only to our marine particle dataset, but any dataset that is subject to distribution shift and a scarcity of labels for minority classes, two challenges which are ubiquitous in ecological image datasets.

## Acknowledgements

This work was supported by the NASA EXORTS program (80NSSC17K0662). VJA was additionally supported by the National Science Foundation Graduate Research Fellowship, the UC Eugene Cota-Robles Fellowship, the Computer Vision for Ecology workshop hosted by the Resnick Sustainability Institute. CAD was additionally supported by the David and Lucile Packard Foundation. We thank the captain and crew of the research vessels on which these data were collected and the science teams who assisted with at-sea operations. We thank Margaret Estapa, Melissa Omand, Ivona Cetinic, and Alyson Santoro, who lead sediment trap deployments during these field campaigns and contributed carbon flux measurements to this study. We also thank Phoebe Lam and Jessica Sheu, who contributed ideas during the early stages of this project.

## Supplemental information

**Figure S1:**
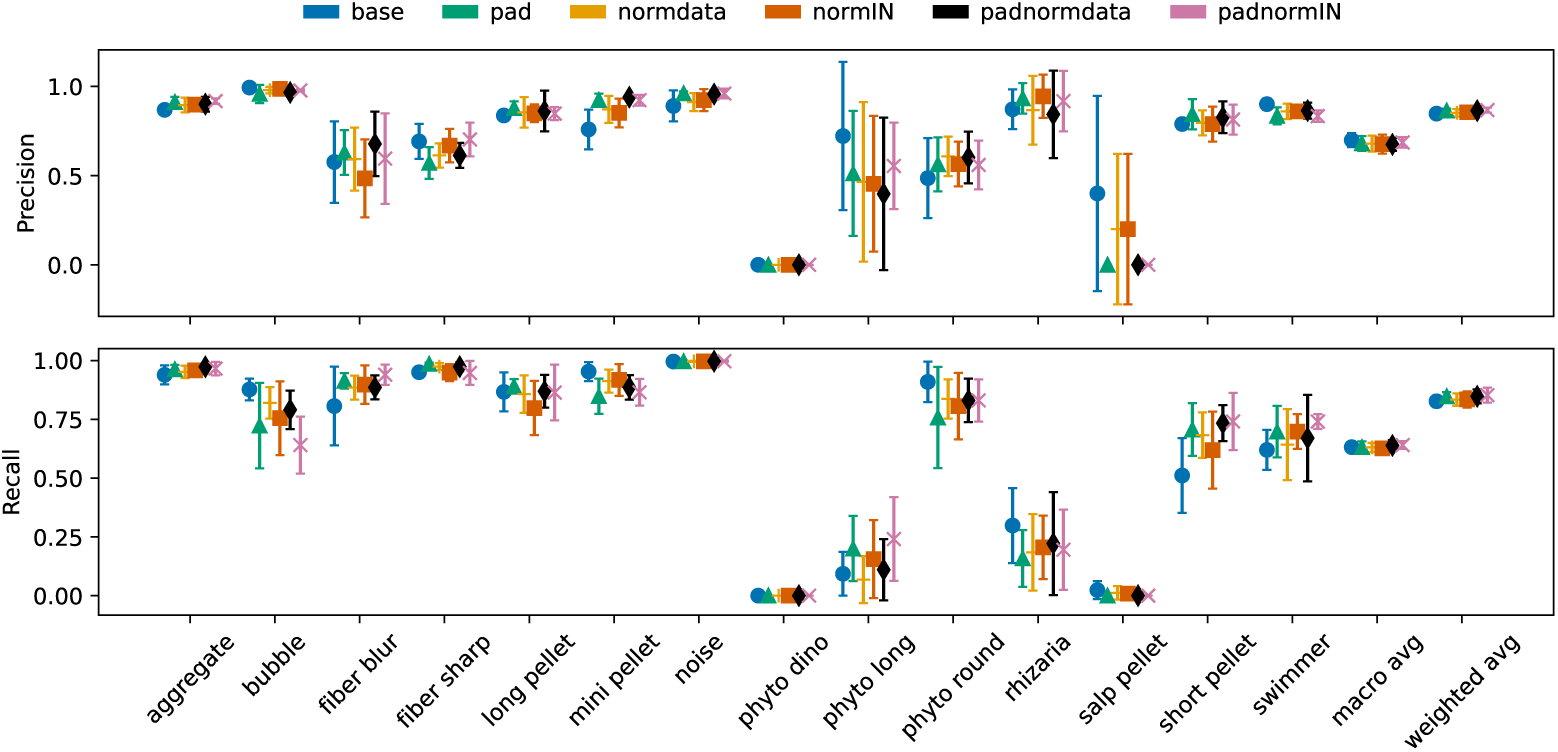
Precision and recall by class for preprocessing protocol tuning. The macro average is the arithmetic mean across all classes, while the weighted average is weighted by the abundance of each class in the total training distribution. Error bars indicate one standard deviation of flux estimates across five model replicates. (base) Resize with no normalization. (pad) CustomPad with no normalization. (normdata) Resize with normalization via statistics calculated from our Resize-transformed data. (normIN) Resize with normalization via ImageNet statistics. (padnormdata) CustomPad with normalization via statistics calculated from our CustomPad-transformed data. (padnormIN) CustomPad with normalization via ImageNet statistics.

**Figure S2:**
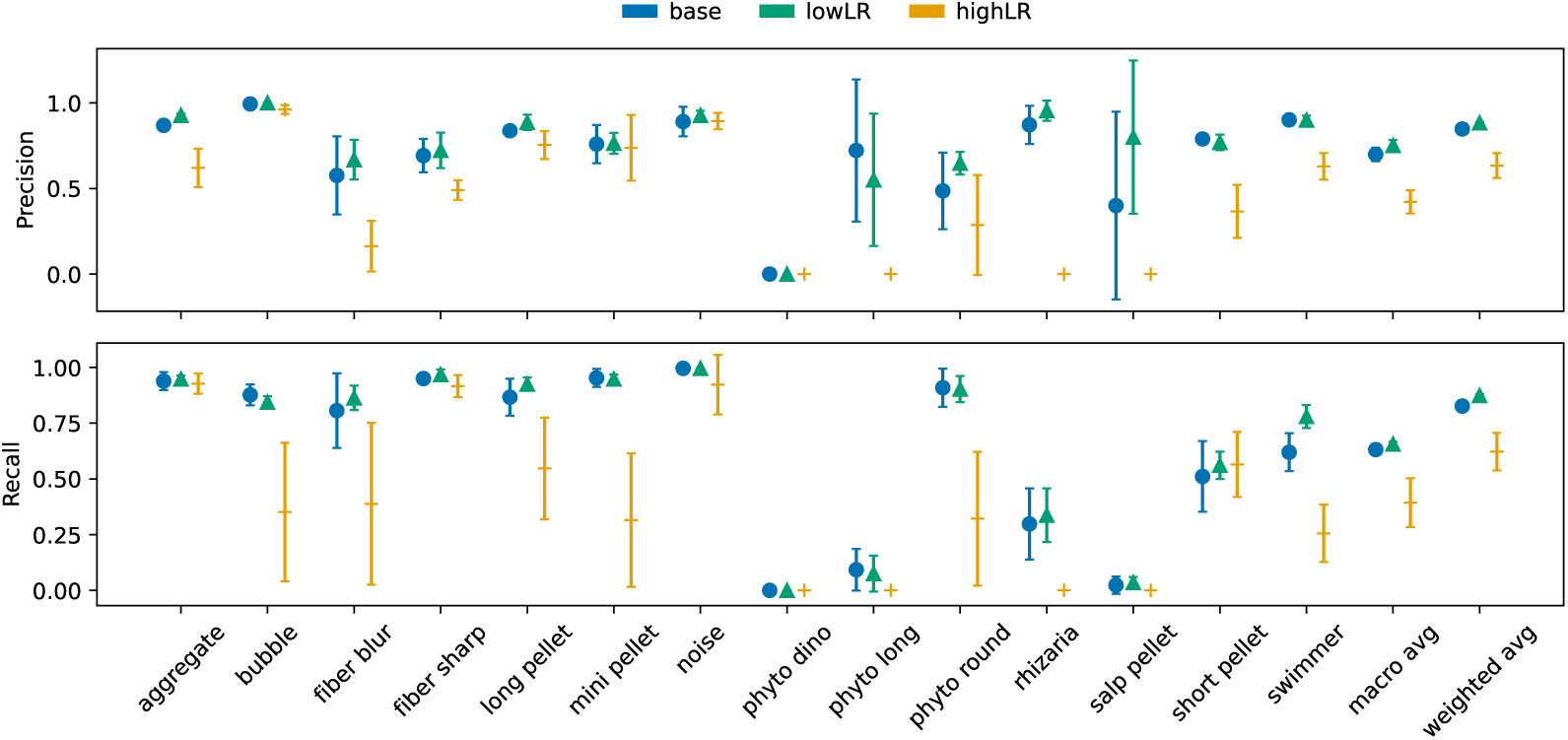
Precision and recall by class for learning rate tuning. The macro average is the arithmetic mean across all classes, while the weighted average is weighted by the abundance of each class in the total training distribution. Error bars indicate one standard deviation of flux estimates across five model replicates. (base) learning rate set to 0.001. (lowLR) 0.0001. (highLR) 0.01.

**Figure S3:**
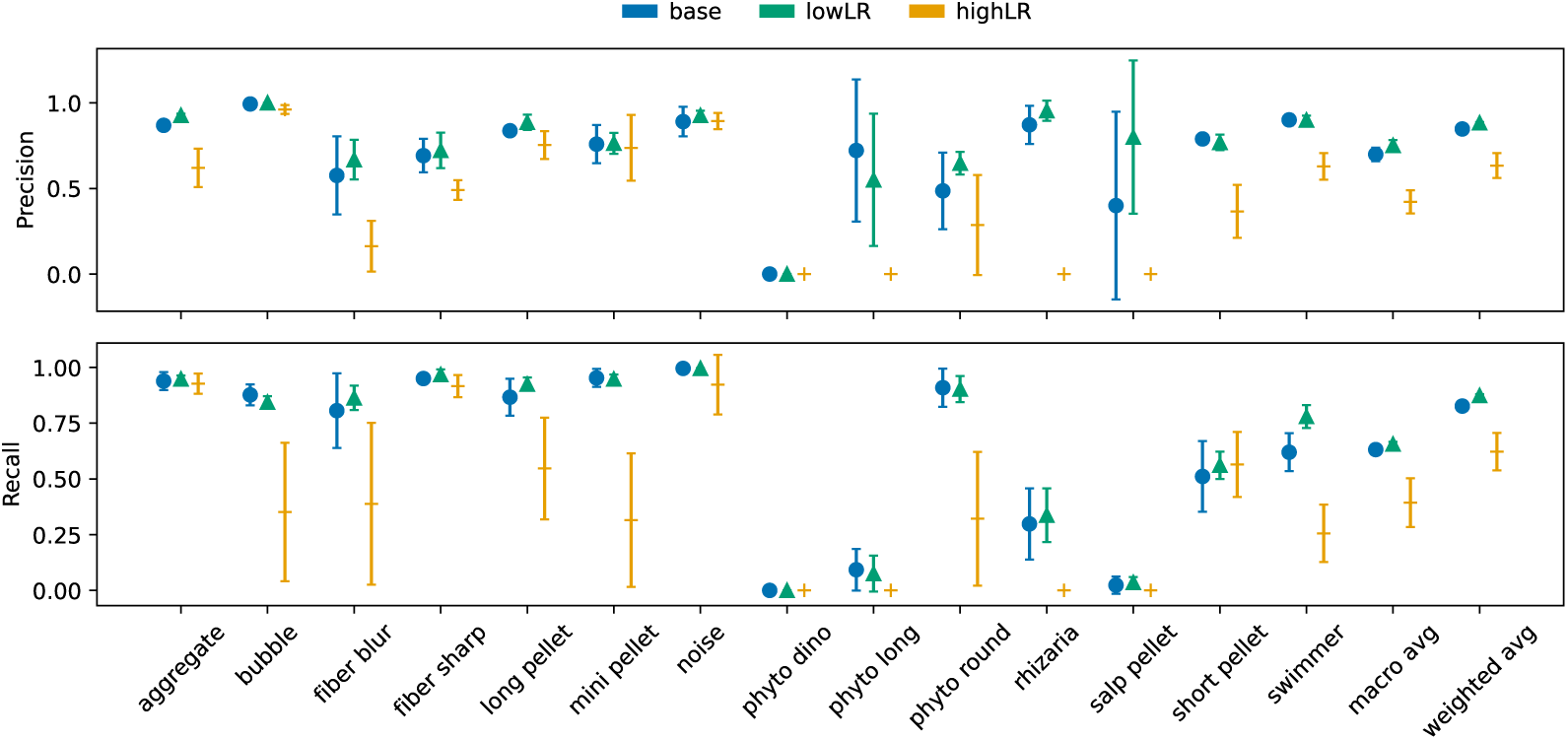
Precision and recall by class for weight decay tuning. The macro average is the arithmetic mean across all classes, while the weighted average is weighted by the abundance of each class in the total training distribution. Error bars indicate one standard deviation of flux estimates across five model replicates. (base) weight decay set to 0.01. (lowWD) 0.001. (highWD) 0.1.

## References

Bisson, K. M., Kiko, R., Siegel, D. A., Guidi, L., Picheral, M., Boss, E., Cael, B. B. 2022. Sampling uncertainties of particle size distributions and derived fluxes. Limnology and Oceanography: Methods, 20(12):754–767.

Boyd, P. W., Claustre, H., Levy, M., Siegel, D. A., Weber, T. 2019. Multi- faceted particle pumps drive carbon sequestration in the ocean. Nature, 568(7752):327–335.

Canziani, A., Paszke, A., Culurciello, E. 2017. An Analysis of Deep Neural Network Models for Practical Applications. arXiv:1605.07678 [cs].

Cheng, K., Cheng, X., Wang, Y., Bi, H., Benfield, M. C. 2019. Enhanced convolutional neural network for plankton identification and enumeration. PLOS ONE, 14(7):e0219570.

Clements, D. J., Yang, S., Weber, T., McDonnell, A. M. P., Kiko, R., Stemmann, L., Bianchi, D. 2022. Constraining the Particle Size Distribution of Large Marine Particles in the Global Ocean With *In Situ* Optical Observations and Supervised Learning. Global Biogeochemical Cycles, 36(5).

Clements, D. J., Yang, S., Weber, T., McDonnell, A. M. P., Kiko, R., Stemmann, L., Bianchi, D. 2023. New Estimate of Organic Carbon Export From Optical Measurements Reveals the Role of Particle Size Distribution and Export Horizon. Global Biogeochemical Cycles, 37(3).

Dai, J., Wang, R., Zheng, H., Ji, G., Qiao, X. 2016. ZooplanktoNet: Deep convolutional network for zooplankton classification. In OCEANS 2016 - Shanghai, pages 1–6, Shanghai, China. IEEE.

Daume III, H., Marcu, D. 2006. Domain Adaptation for Statistical Classifiers. Journal of Artificial Intelligence Research, 26:101–126.

Ducklow, H., Steinberg, D., Buesseler, K. 2001. Upper ocean carbon export and the biological pump. Oceanography, 14(4):50–58.

Durkin, C. A., Buesseler, K. O., Cetinić, I., Estapa, M. L., Kelly, R. P., Omand, M. 2021. A visual tour of carbon export by sinking particles. Global Biogeochemical Cycles, 35(10).

Durkin, C. A., Estapa, M. L., Buesseler, K. O. 2015. Observations of carbon export by small sinking particles in the upper mesopelagic. Marine Chemistry, 175:72–81.

Estapa, M., Buesseler, K., Durkin, C. A., Omand, M., Benitez-Nelson, C. R., Roca-Martí, M., Breves, E., Kelly, R. P., Pike, S. 2021. Biogenic sinking particle fluxes and sediment trap collection efficiency at Ocean Station Papa. Elementa: Science of the Anthropocene, 9(1).

Giering, S. L. C., Cavan, E. L., Basedow, S. L. and others. 2020. Sinking organic particles in the ocean—Flux estimates from in situ optical devices. Frontiers in Marine Science, 6.

Guo, B., Nyman, L., Nayak, A. R., Milmore, D., McFarland, M., Twardowski, M. S., Sullivan, J. M., Yu, J., Hong, J. 2021. Automated plankton classification from holographic imagery with deep convolutional neural networks. Limnology and Oceanography: Methods, 19(1):21–36.

Guo, C., Pleiss, G., Sun, Y., Weinberger, K. Q. 2017. On Calibration of Modern Neural Networks. In Proceedings of the 34 th International Conference on Machine Learning, Sydney, Australia.

Hashemi, M. 2019. Enlarging smaller images before inputting into convolutional neural network: zero-padding vs. interpolation. Journal of Big Data, 6(1):98.

He, K., Zhang, X., Ren, S., Sun, J. 2016. Deep Residual Learning for Image Recognition. In 2016 IEEE Conference on Computer Vision and Pattern Recognition (CVPR), pages 770–778, Las Vegas, NV, USA. IEEE.

Hong, S., Raza, S., Huang, H., Shahani, K., Zhang, Y.,, J., Raza, K., Ali, M. 2020. Classification of Freshwater Zooplankton by Pre-trained Convolutional Neural Network in Underwater Microscopy. International Journal of Advanced Computer Science and Applications, 11(7).

Iversen, M. H., Pakhomov, E. A., Hunt, B. P., Van Der Jagt, H., Wolf-Gladrow, D., Klaas, C. 2017. Sinkers or floaters? Contribution from salp pellets to the export flux during a large bloom event in the Southern Ocean. Deep Sea Research Part II: Topical Studies in Oceanography, 138:116–125.

Johnson, L., Siegel, D. A., Thompson, A. F. and others. 2024. Assessment of oceanographic conditions during the North Atlantic EXport processes in the ocean from RemoTe sensing (EXPORTS) field campaign. Progress in Oceanography, 220:103170.

Kay, J., Kulits, P., Stathatos, S., Deng, S., Young, E., Beery, S., Van Horn, G., Perona, P. 2022. The Caltech Fish Counting Dataset: A Benchmark for Multiple-Object Tracking and Counting. In Avidan, S., Brostow, G., Cissé, M., Farinella, G. M., Hassner, T., editors, Computer Vision – ECCV 2022, volume 13668, pages 290–311. Springer Nature Switzerland, Cham. Series Title: Lecture Notes in Computer Science.

Li, Y., Guo, J., Guo, X., Zhao, J., Yang, Y., Hu, Z., Jin, W., Tian, Y. 2021. Toward in situ zooplankton detection with a densely connected YOLOV3 model. Applied Ocean Research, 114:102783.

Lombard, F., Boss, E., Waite, A. M. and others. 2019. Globally Consistent Quantitative Observations of Planktonic Ecosystems. Frontiers in Marine Science, 6:196.

Loshchilov, I., Hutter, F. 2019. Decoupled Weight Decay Regularization.

Menden-Deuer, S., Lessard, E. J. 2000. Carbon to volume relationships for dinoflagellates, diatoms, and other protist plankton. Limnology and Oceanography, 45(3):569–579.

Orenstein, E. C., Beijbom, O. 2017. Transfer Learning and Deep Feature Extraction for Planktonic Image Data Sets. In 2017 IEEE Winter Conference on Applications of Computer Vision (WACV), pages 1082–1088, Santa Rosa, CA, USA. IEEE.

Orenstein, E. C., Kenitz, K. M., Roberts, P. L., Franks, P. J., Jaffe, J. S., Barton, A. D. 2020. Semi- and fully supervised quantification techniques to improve population estimates from machine classifiers. Limnology and Oceanography: Methods, 18(12):739–753.

Picheral, M., Guidi, L., Stemmann, L., Karl, D. M., Iddaoud, G., Gorsky, G. 2010. The Underwater Vision Profiler 5: An advanced instrument for high spatial resolution studies of particle size spectra and zooplankton: Underwater vision profiler. Limnology and Oceanography: Methods, 8(9):462–473.

Russakovsky, O., Deng, J., Su, H. and others. 2015. ImageNet Large Scale Visual Recognition Challenge. International Journal of Computer Vision, 115(3):211–252.

Schröder, S.-M., Kiko, R. 2022. Assessing Representation Learning and Clustering Algorithms for Computer-Assisted Image Annotation—Simulating and Benchmarking MorphoCluster. Sensors, 22(7):2775.

Schröder, S.-M., Kiko, R., Koch, R. 2020. MorphoCluster: Efficient Annotation of Plankton Images by Clustering. Sensors, 20(11):3060.

Siegel, D. A., Cetinić, I., Graff, J. R. and others. 2021. An operational overview of the EXport Processes in the Ocean from RemoTe Sensing (EXPORTS) Northeast Pacific field deployment. Elementa: Science of the Anthropocene, 9(1).

Silver, M. W., Bruland, K. W. 1981. Differential Feeding and Fecal Pellet Composition of Salps and Pteropods, and the Possible Origin of the Deep- Water Flora and Olive-Green ”Cells”. Marine Biology, 62:263–273.

Stukel, M. R., Biard, T., Krause, J., Ohman, M. D. 2018. Large Phaeodaria in the twilight zone: Their role in the carbon cycle. Limnology and Oceanography, 63(6):2579–2594.

Trudnowska, E., Lacour, L., Ardyna, M., Rogge, A., Irisson, J. O., Waite, A. M., Babin, M., Stemmann, L. 2021. Marine snow morphology illuminates the evolution of phytoplankton blooms and determines their subsequent vertical export. Nature Communications, 12(1):2816.

Zhou, K., Liu, Z., Qiao, Y., Xiang, T., Loy, C. C. 2022. Domain Generalization: A Survey. IEEE Transactions on Pattern Analysis and Machine Intelligence, pages 1–20.

